# Nasal respiration entrains neocortical long-range gamma coherence during wakefulness

**DOI:** 10.1101/430579

**Authors:** Matías Cavelli, Santiago Castro-Zaballa, Joaquín Gonzalez, Daniel Rojas-Líbano, Nicolas Rubido, Noelia Velásquez, Pablo Torterolo

## Abstract

Recent studies have shown that slow cortical potentials in archi-, paleo- and neocortex, can phase-lock with nasal respiration. In some of these areas, gamma activity (γ: 30-100 Hz) is also coupled to the animal’s respiration. It has been hypothesized that this interaction plays a role in coordinating distributed neural activity. In a similar way, inter-cortical interactions at γ frequency has been also associated as a binding mechanism by which the brain generates temporary opportunities necessary for implementing cognitive functions. The aim of the present study is to explore if nasal respiration entrains inter-cortical interactions at γ frequency.

Six adult cats chronically prepared for electrographic recordings were employed in this study. Our results show that slow cortical respiratory potentials are present in several areas of the neocortex and olfactory bulb during wakefulness. Also, we found cross-frequency coupling between the respiratory phase and the amplitude of γ activity in all recorded areas. These oscillatory entrainments are independent of muscular activity, because are maintained during cataplexy induced by carbachol microinjection into the nucleus pontis oralis. Importantly, we observed that respiratory phase modulates the inter-cortical gamma coherence between neocortical pairs of electrodes during wakefulness. However, during NREM and REM sleep, breathing was unable to entrain the oscillatory activity, neither in the olfactory bulb nor in the neocortex. These results suggest a single unified phenomenon involving cross frequency coupling and long-range γ coherence across the neocortex. This fact could be related to a temporal binding process necessary for cognitive functions during wakefulness.

## Introduction

The brain is a complex system, in which parallel processing coexist with serial operations within highly interconnected networks, but without a single coordinating center. This organ integrates neural events that occur at different times and locations into a unified perceptual experience. Understanding the mechanisms responsible for this integration, is a crucial challenge for cognitive neuroscience^1–3^

Neural synchronization at gamma frequency band (γ: 30-100 Hz) is hypothesized as a binding mechanism through which the brain generates transient opportunities for communication and integration of the distributed neural activity necessary for cognitive functions^1, 3–7^. Cortical γ power increases during active behavioral states as well as during the performance of cognitive tasks^2, 5, 8–11^. Besides power, γ synchronization between distant areas of the brain (γ coherence) also increases during several cognitive functions in both animals and humans^2, 5, 9, 12, 13^. γ coherence has been been considered as a neural correlate of conscious perception^5, 13–15^, it decreases during sleep^2, 9^ and is absent during narcosis (unconsciousness) induced by general anesthetics^16, 17^. Recently, it was shown that slow oscillations such as the theta rhythm of the hippocampal networks^18,19^, the cortical potentials caused by the rhythmic movement of the eyes^20,21^ and respiration^19, 22, 23^, can modulate γ activity.

The work of Adrian^24^ is usually regarded as the first description of nasal respiration driving neural oscillations in the olfactory bulb (OB). In the nasal epithelium, inhaled air activates olfactory sensory neurons (OSNs), which can detect both odor and mechanical stimuli^25,26^. Air flowing through the nostrils can synchronize the neuronal activity and local field potentials (LFPs) in OB and piriform cortex^11,27^. Recently, it has been observed that breathing also couples the slow activity of other brain areas not directly related to olfaction. For example, Ito et al.^23^ showed that spikes and delta oscillations from the somatosensory cortex can phase-lock with respiration in awake mice. This locking or cortical respiratory potential (CRP) is lost after bulbectomy. CRP were also observed in the dorsal hippocampus of rats and mice, prominently in the dentate gyrus^28,29^, in medial and orbitofrontal prefrontal cortex (Pf)^19,30,31^, primary visual (V1) and primary motor (M1) cortex^32^. Other studies have showed that CRP also occurs in several regions of the human brain^33,34^. Researchers have also observed respiratory modulation of local γ activity in most of the abovementioned areas 10, 11, 19, 23, 29, 30, 32. This “Cross Frequency Coupling” (CFC)^35^ has been also hypothesized by some authors to play a potential role in integrating distributed network activity^19,22^.

Since this modulation of brain waves depends on air passage through the nostrils, and on an intact OB, it is probably a breathing reafferent signal that modulates multiples areas of the brain^22,32^.

Employing the cat as an animal model, the aim of the present study was to seek if slow regional oscillatory activity phase-locks to respiration, and couples with the γ activity in cortical areas during wakefulness (W) and sleep. In addition, we studied if nasal respiration modulates inter-cortical long-range γ coherence, especially between areas that were not previously related to the sensorimotor act of breathing.

## Results

### Cortical respiratory potentials are present during wakefulness but not during sleep

Electrocorticogram recordings (ECoG) from several areas of the neocortex and OB (Figure S1) were obtained during the natural sleep-wake cycle of six cats. We simultaneously monitored the nuchal electromyogram (EMG), and the respiratory activity through a thermistor in the nostrils and a micro-effort sensor in the chest. Recordings were performed in a head-restricted condition with the body resting in a sleeping bag^9, 36, 37^.

First, we determined the presence of CRP in the ECoG, and its dependence on the animal’s behavioral state. Figure 1A shows the polysomnographic recording of a representative cat during W, non-REM (NREM) sleep and REM sleep. During W, we observed that slow respiratory waves were accompanied by high amplitude oscillations of similar frequency in the OB; similar potentials of lower amplitude were also present in the neocortex. During NREM sleep, although we observed the characteristic slow waves and sleep spindles, these oscillations do not seem associated with the respiratory cycle. During REM sleep, CPRs were not observed in any of the recorded areas (Figure 1A).

**Figure 1.**
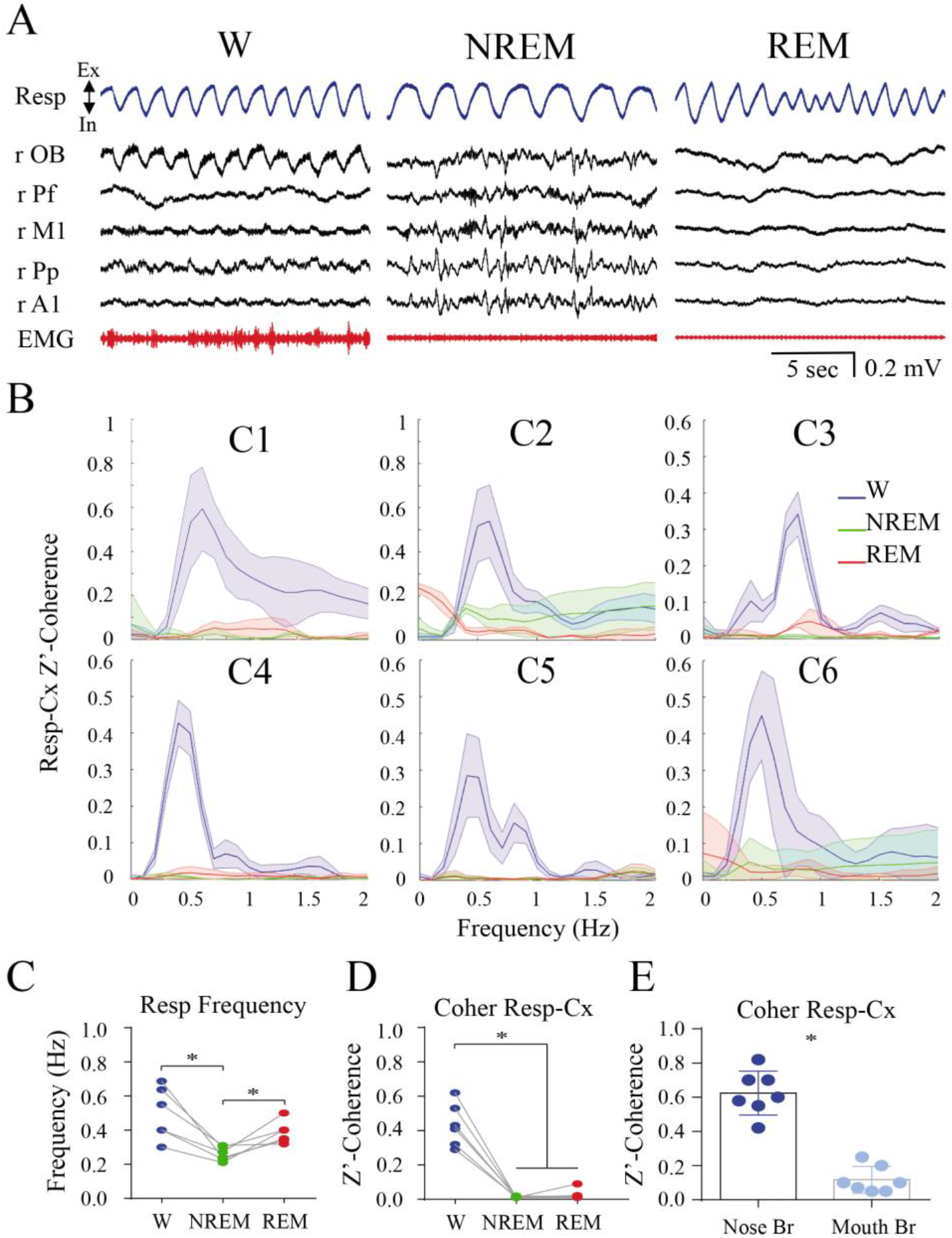
Cortical respiratory potential (CRP) in the cat’s cortex during wakefulness (W) and sleep. A. Simultaneous polysomnographic recordings during W, NREM and REM sleep from a representative animal (C1). Breathing was recorded through a thermistor in the nostrils (blue) in simultaneous with the ECoG (black). The recordings are from the olfactory bulb (OB), prefrontal cortex (Pf), primary motor cortex (M1), posterior parietal cortex (Pp) and primary auditory cortex (A1) from the right hemisphere (r). The EMG recordings are shown in red. In, inhalation; Ex, exhalation. B. Magnitude squared coherence between the respiratory waves and ECoG signals during sleep and W. We applied the Fisher z-transform to the coherence values (z’-coherence). Each graph represents the mean ± standard deviation of all the pairs between the respiratory waves and each ECoG area recording for each cat (C1-C6; see Figure S1 for the electrode location in each animal) during W, NREM and REM sleep. C. Respiratory frequency during W and sleep. The respiratory frequency was extracted from the power spectrum peak of the respiratory signal. The statistical significance was analyzed with repeated measures ANOVA and Bonferroni *post hoc* tests (* p<0.05). D. Maximal mean z’-coherence between respiratory wave and ECoG, calculated at the respiratory frequency during W, NREM and REM sleep. The statistical significance is the same as that used in C. E. Z’-coherence between respiratory wave and ECoG at the respiratory frequency during mouth and nose breathing. The graph represents the mean ± standard deviation of seven cortical areas of one animal (C2). The coherence analysis was performed on ten concatenated segments of 50 seconds (500 sec), where the animal was forced to breathe through the mouth or freely through the nose. The statistical significance was evaluated by means of the two-tailed paired t-test. (* p<0.05). Br, breathing.

We also found in all the animals that respiration and cortical activity were spectrally coherent during W, but not during NREM and REM sleep (Figure 1B). We then calculated the respiratory rate through spectral analysis of the respiratory signal. As expected, the respiratory frequency was dependent of the behavioral state (rmANOVA; F_(1.144, 5.721)_=8.158; p=0.028). During NREM sleep, respiration rate was lower in comparison to W and REM sleep (Figure 1A and C)^37^. Thereafter, we computed the average coherence levels for each animal at the peak of the respiratory wave power spectrum^22^ Figure 1D shows that coherence values between respiratory oscillations and ECoG are large during W and fall during sleep (rmANOVA; F_(1.094, 5.469)_=66.56; p=0.0003). Next, we sought to determine whether these CRPs were related to the passage of air through the nostrils. As shown in Figure 1E, the coherence between respiration and ECoG that is observed during nasal respiration in W, is lost during mouth breathing.

Transitions from sleep to W are shown in Figure 2A; CRPs are clearly associated with W.

**Figure 2.**
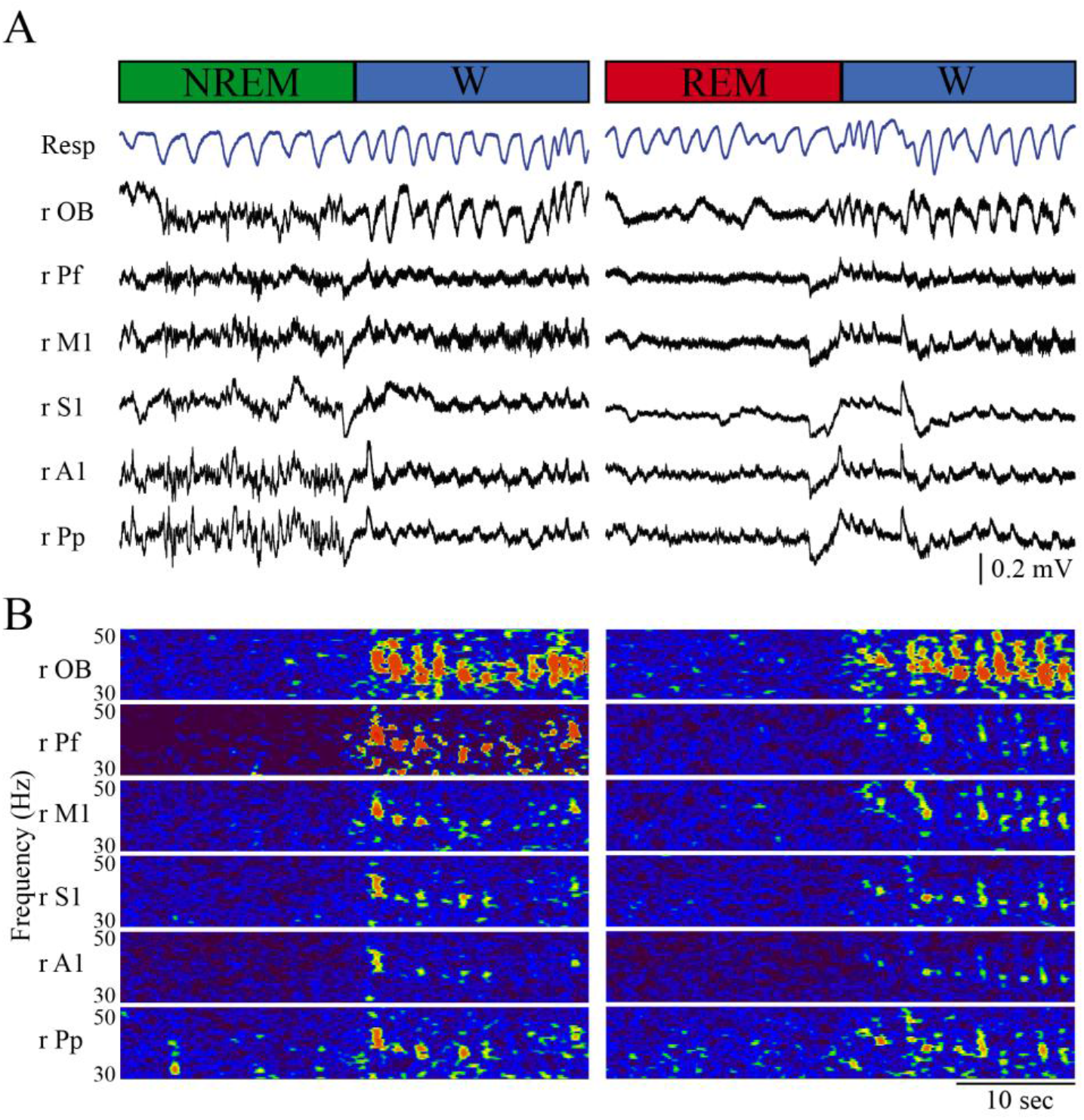
Cortical respiratory potential (CRP) during sleep to wakefulness (W) transitions. A. Simultaneous polysomnographic recordings during NREM sleep to W (left), and REM sleep to W (right) transitions (C1). Breathing was recorded with a thermistor in the nostrils (blue) in simultaneous with the ECoG (black). The recordings are from the olfactory bulb (OB), prefrontal cortex (Pf), primary motor cortex (M1), primary somatosensory cortex (S1), posterior parietal cortex (Pp) and primary auditory cortex (A1) from the right hemisphere (r). B. Time-frequency power decomposition during behavioral states transitions. 30-50 Hz power spectrogram (1 second and 0.5 Hz resolution).

### Respiration entrains cortical γ activity during wakefulness

Spectrograms of Figure 2B and filtered recordings shown in Figure S2, exhibit that bursts of coupled γ band (30-50 Hz) activity are also related with W and associated with breathing. We computed CFC between respiration and γ oscillations (CFC(Resp-γ)) through cross-correlation function maps (CCFmap)^18^ between the respiratory signal and the amplitude envelopes of 10 to 100 Hz oscillations. Figure 3 shows the CCFmap of 5 neocortical areas and the OB of a representative animal during W, NREM and REM sleep, as well as auto-correlation function (ACF) of the respiratory wave. We observed a clear cross-correlation between respiration and γ activity for all the recorded areas during W. On the other hand, during sleep (NREM and REM) the CFC(Resp-γ) levels are negligible. In the ACF, the value of zero corresponds to the end of expiration and beginning of inspiration. We can see that the γ-respiration correlation increases mainly during the expiratory phase of the cycle. On the other hand, the end of inspiration is accompanied by higher gamma frequencies that become progressively lower as expiration develops (Figure 3, W, 0 Lag). We found similar behavioral state dependent changes in CFC(Resp-γ) for each recorded animal (see Figure S3). This analysis also reveals some variability among the animals regarding the frequency limits of the γ burst (Figure S3); in some animals the frequency range of the bursts goes up to 60 Hz.

**Figure 3.**
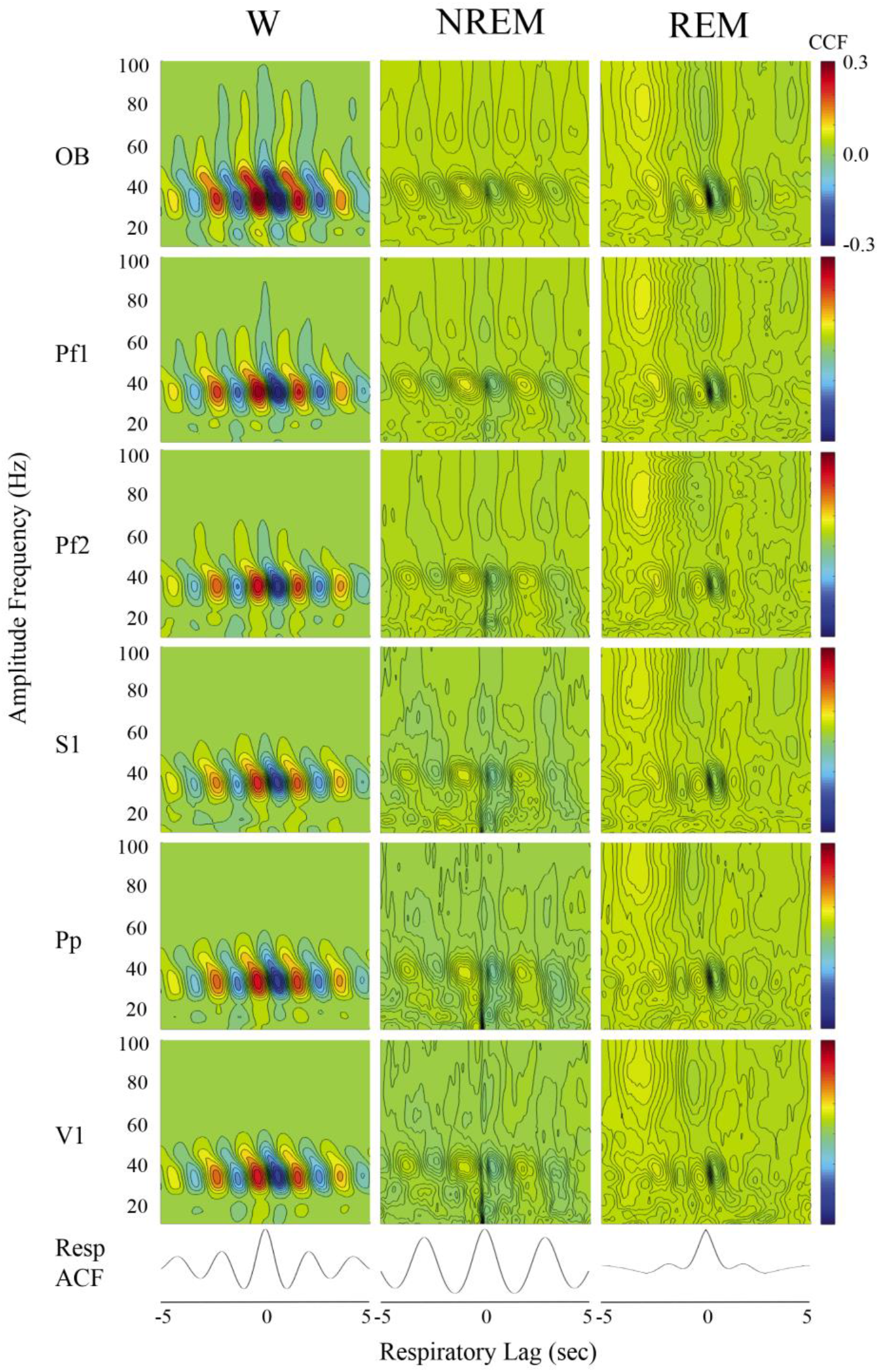
Top: CFC(Resp-γ) during wakefulness and sleep. Cross-correlation function (CCF) maps between the respiratory wave and the amplitude envelope of the ECoG signals between 10 and 100 Hz (10-Hz bandwidth and 5-Hz steps) during wakefulness (W), NREM, and REM sleep. Bottom: autocorrelation function (ACF) of the respiratory signal. The CCF maps correspond to the olfactory bulb (OB), anterior prefrontal cortex (Pf1), posterior prefrontal cortex (Pf2), primary somatosensory cortex (S1), posterior parietal cortex (Pp) and primary visual cortex (V1) of the right hemisphere (C2; see Figure S1 for electrode position).

Previous reports have found an increase in the respiratory frequency variability especially during REM sleep^37^. This is reflected in the loss of the periodic auto-correlation observed during REM sleep (Figure 3, ACF)^37^, and could be the cause of the low levels of cross-correlation observed during this state. In order to discard that the decrease in the CFC(Resp-γ) levels during REM sleep is caused by this increment in respiratory variability^37^, and due to the importance of phase coding in olfactory perception^25^, we analyzed the relationships between the phase (in degrees) of the respiratory wave and the amplitude (envelopes) of γ activity (phase amplitude coupling or PAC), using the modulation index (MI) designed by Tort *et al*^35^. The phase-amplitude MI quantifies the deviation of the empirical phase-amplitude distribution from a uniform distribution. Figure 4A shows the average MI(Resp-γ) values of all the areas recorded for each animal (n=6) during W and sleep. This analysis revealed that the highest MI(Resp-γ) values were observed during W (Figure 4A; rmANOVA; F_(1.011, 5.057)_=14.45; p=0.0123). Then, we analyzed how the MI(Resp-γ) values varied depending on the type of breathing, buccal or nasal. Figure 4B shows the MI value for all areas of a representative animal recorded during buccal and nasal breathing during W; MI(Resp-γ) was significantly higher during nasal breathing.

**Figure 4.**
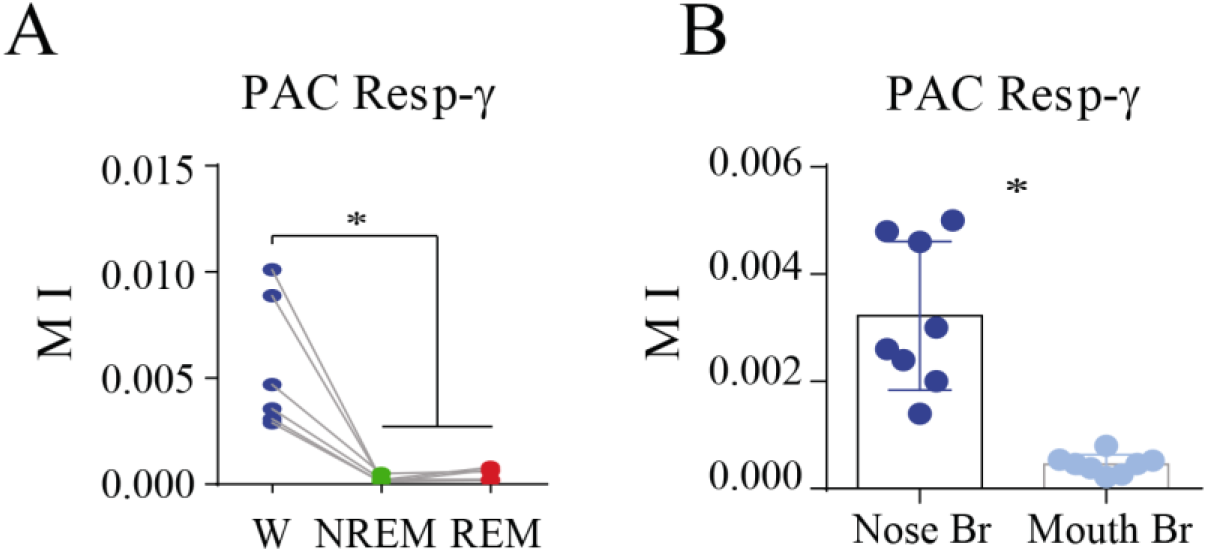
PAC(Resp-γ) during wakefulness (W), NREM and REM sleep. A. Mean phase amplitude coupling between the phase of the respiratory waveform and the amplitude (envelope) of the γ activity, computed using the modulation index (MI) as a measure of the CFC level between the two signals^35^. Each value represents the mean MI for all the areas recorded for each animal (n = 6). The statistical significance among behavioral states was evaluated with repeated measures ANOVA and Bonferroni *post hoc* tests (* p<0.05). B. MI between respiratory phase and the γ envelopes during mouth and nose breathing. The graph represents the mean ± standard deviation of seven cortical areas of one animal (C1). The MI analysis was performed on ten concatenated segments of 50 seconds (500 sec) where the animal was forced to breathe through the mouth or freely through the nose. The statistical significance was evaluated with the two-tailed paired t-test (* p<0.05). Br, breathing; PAC, phase amplitude coupling.

### Respiratory cortical entrainment does not depend on muscular tone

One of the main differences between W and sleep is muscle tone^2,9,36,37^, which is the main artefactual signal recorded in the EEG, ECoG and LFPs during W^8^. To rule out the possibility that muscle activity was affecting the results, we carried out experiments where we turned off the muscular activity. Microinjections of carbachol (cholinergic agonist) in the nucleus pontis oralis (NPO) causes the animal to alternate between states of cataplexy (CA; W with atony) and REM sleep induced by carbachol (REMc; also, with atony) for 30 minutes to 2 hours. Upon carbachol microinjections, we found that although the animals exhibited muscle atonia in both states (CA and REMc), only during CA we observed CRP (Figure 5A and B), and an increase in CFC(Resp-γ) (Figure 5C and D) between respiratory oscillations and ECoG.

**Figure 5.**
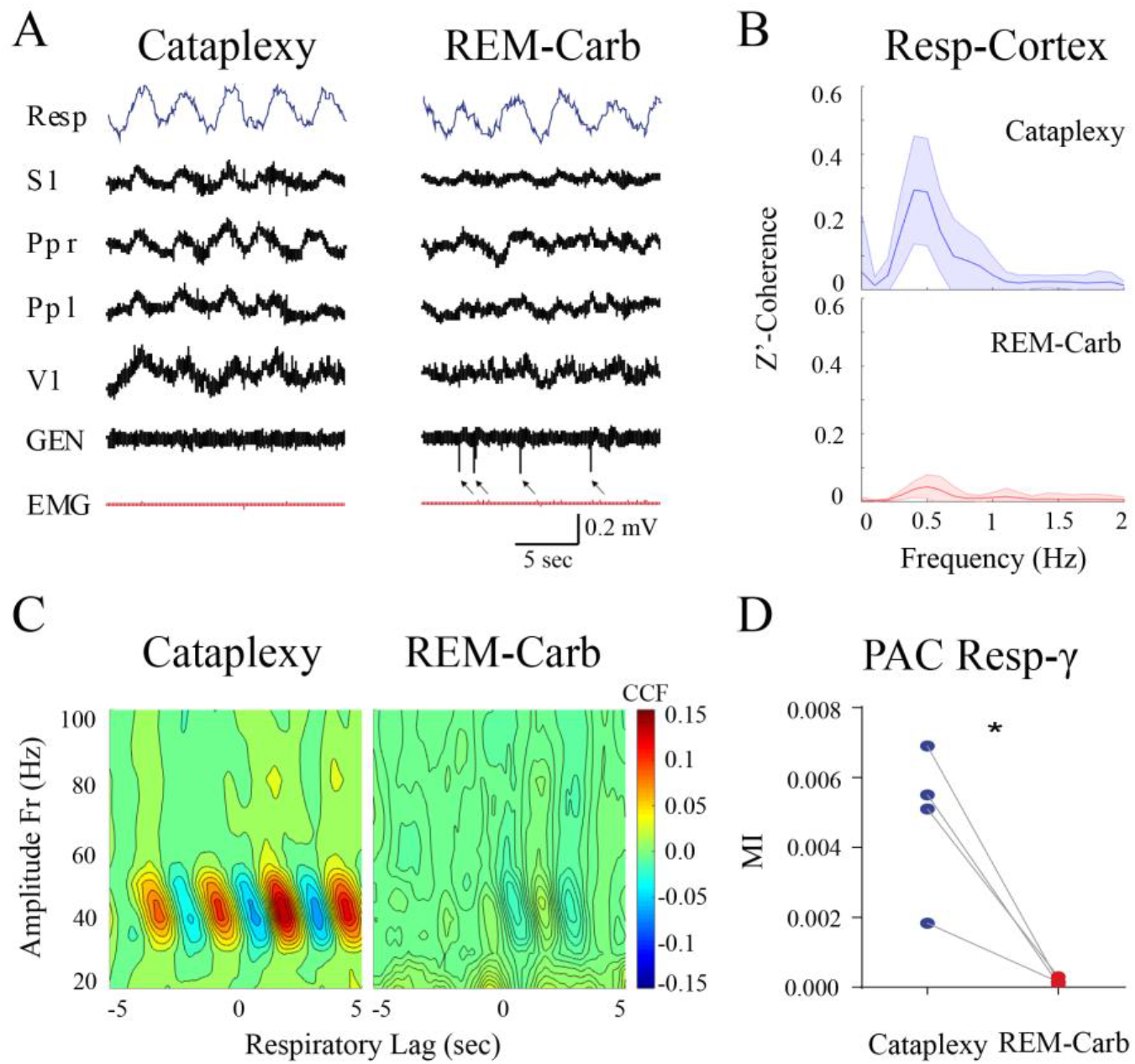
CRP and CFC(Resp-γ) were independent of the muscular tone. A. Simultaneous polysomnographic recordings during cataplexy (CA) and REM sleep induced by carbachol (REMc). Breathing was recorded with a micro-effort piezo crystal infant sensor in simultaneous with the ECoG (black) from the primary sensory cortex (S1), left and right posterior parietal cortex (Pp l and Pp r) and primary visual cortex (V1). Lateral geniculate nucleus (LGN) electrogram and EMG are also shown (red). Muscle tone absence is observed in both states, but only during REMc PGO waves were present in the LGN (arrows). B. Z’-coherence between the respiratory wave and ECoG signals during CA and REMc. Each graph represents the mean ± standard deviation of all the cortical areas, and all the animals (n = 4) (Figure S1 for the electrode location in each animal; C3-C6). C. Mean CCF maps during CA and REMc. Mean cross-correlation function (CCF) between the respiratory waves and the amplitude envelope of the ECoG signals (between 10 and 100 Hz; 10-Hz bandwidth and 5-Hz steps) during CA, and REMc. The CCF map for each condition corresponds to the mean of all the CCF maps of every area and animal. D. PAC(Resp-γ) during CA and REMc. Each value represents the mean MI value for all areas of each animal. The statistical significance between CA and REMc was evaluated with the two-tailed paired t-test (* p<0.05).

### Respiration entrains neocortical long-range γ coherence

To investigate if respiratory rhythms facilitate inter-cortical communication through high frequency channels, we studied the co-modulation between the respiratory waves and the cortico-cortical γ synchronization using different metrics. Figure 6A shows a polysomnographic raw recording of a representative animal during W. The same recordings filtered for the γ band (30-60 Hz; blue) and superimposed the amplitudes (RMS envelopes) of the same filtered records (red) are exhibited in Figure 6B; it is readily observed that γ amplitude seems to be in phase with the respiratory wave. In addition, we analyzed the coherence of all pairs of cortices recorded as a function of the respiratory phase. Figure 6C shows the differences in coherence among behavioral states for all the animals and pairs of electrodes recorded. During W, γ coherence modulates the respiratory phase and this phenomenon is absent during sleep. We obtained similar results when we analyzed the normalized phase locking value (PLV) (Figure S4). Figure 6D shows that gamma coherence between all the neocortical recording sites was phase-locked with the respiratory signal. However, when coherence was analyzed between neocortical areas and OB, there was not a clear phase preference (Figure 6D). In fact, γ coherence is mostly present between neocortical pair of electrodes (Figure 6E). In contrast, OB-including pairs do not exhibit this phenomenon, despite each of the recorded areas showing CFC(Resp-γ) (Figure 3, 4 and S3).

**Figure 6.**
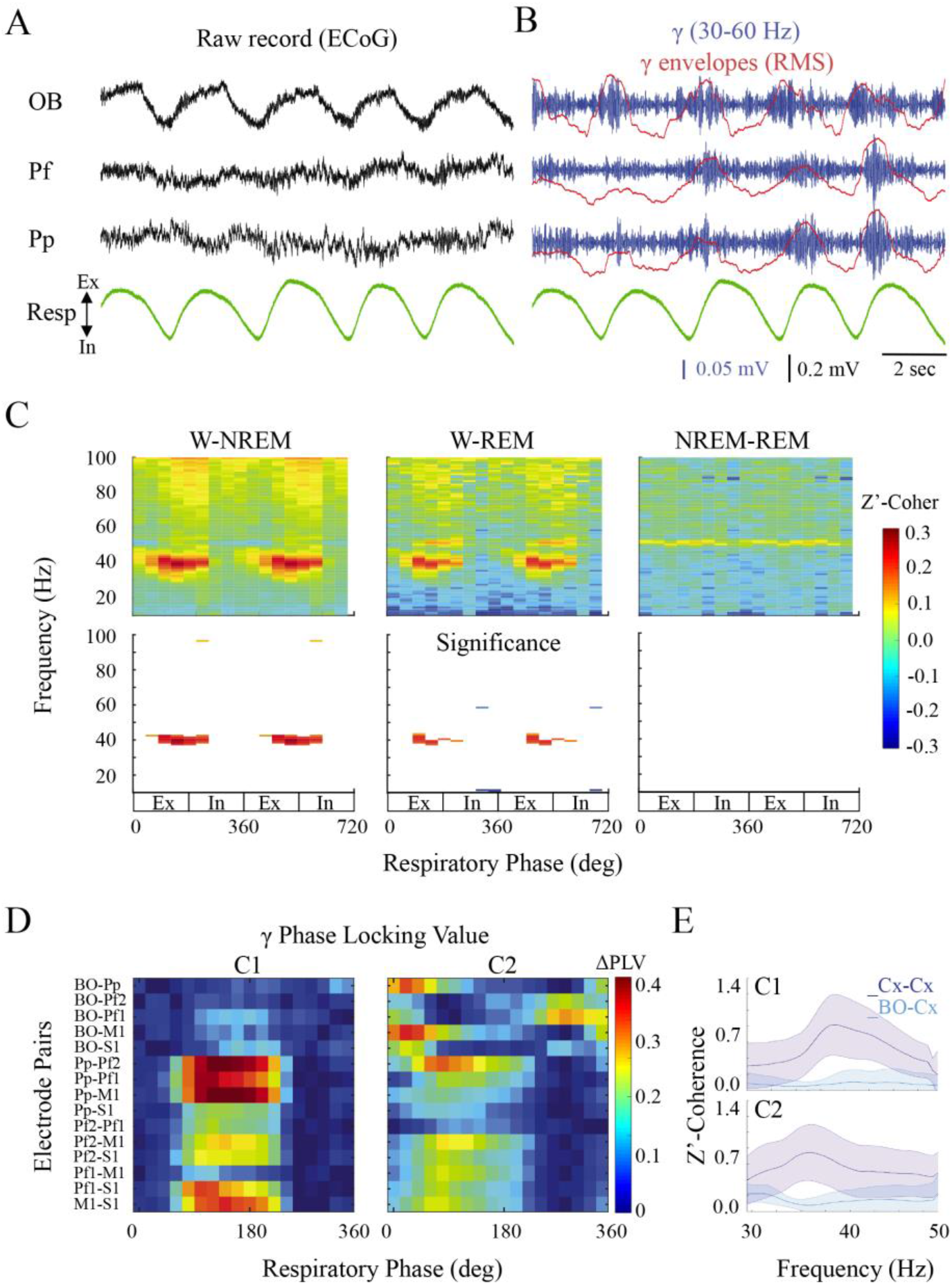
Respiratory modulation of inter-cortical γ coherence. A. The figure exhibits the simultaneous respiratory and ECoG recordings from the olfactory bulb (OB), pre-frontal (Pf), posterior parietal cortex during wakefulness (W). In, inhalation; Ex, exhalation. B. The same ECoG recordings filtered for the gamma band (blue) and its envelope (root mean square, RMS; red) are shown. C. Inter-cortical coherence as a function of the respiratory phase. The respiratory phase was binned 9 times (40 degrees each) and the z’-coherence was calculated for each respiratory bin with a frequency resolution of 0.5 Hz. Each graph represents the mean difference between two behavioral states (W-NREM, W-REM and NREM-REM sleep) of all the animals, and all the ECoG recordings pairs. Below, only the values that were statistically significant in the paired comparison are shown (paired t-test, p < 0.0001). D. γ phase coherence as a function of the respiratory phase for each cortical pair of electrodes is shown. Phase locking values (PLV) was calculated for each specific γ band (30-60 Hz) pair, and then the mean PLV was extracted for each respiratory phase bin (18 degrees per bin). This analysis was done in C1 and C2, because these animals shared the greater number of electrodes in the same cortical areas and OB. To obtain only the values of PLV effectively modulated by respiration, each PLV is subtracted from the minimum found throughout the respiratory cycle (ΔPLV). OB, Olfactory bulb; Pf1, anterior prefrontal cortex, Pf2, posterior prefrontal cortex; M1, primary motor cortex; Pp, posterior parietal cortex; and S1, primary somatosensorial cortex, from the right hemisphere. E. Inter cortical γ z’-coherence between the neocortical pairs (Cx-Cx) and between the OB and the neocortical pairs (OB-Cx). In, inhalation; Ex, exhalation.

The relationship between the gamma synchronization with the respiratory phase was also determined by the analysis of the coherence between the coefficient of determination (R^2^) from all pairs of filtered recordings (γ: 30-60 Hz) and the respiratory signal (Figure S5A); a large coherence was observed only during W. Moreover, R ^2^ values were modulated by the respiratory phase, especially during W (Figure S5B).

## Discussion

Our findings provide evidence about the ability of nasal respiration to entrain neural oscillatory activity in several regions of the cat’s brain. We found that nasal respiration modulates the slow CRP and CFC(Resp-γ) in all recorded areas, in a behavioral state-dependent fashion. Furthermore, we demonstrated that the phase of the respiratory cycle modulates inter-cortical γ coherence during W. This fact could suggest that cross-frequency modulation between respiration and cortical γ rhythms on one side, and long-range inter-cortical γ coherence on the other, could be components of a single phenomenon.

### Cortical respiratory potentials

The existence of CRP in the OB is a well-known phenomenon^10,11,24^. It has been shown that the slow activity of the OB faithfully follows respiration in freely behaving rats^11^; moreover, respiratory slow potentials can be recorded in olfactory and non-olfactory cortical areas^27,19,23,29,32,38^. In the present work, we demonstrated the presence of CRPs in the neocortex of the cat.

### Cross frequency coupling between respiration and γ activity

CFC has been reported in electrophysiological signals such as membrane potential, LFPs, ECoG and EEG^35^. The best-known CFC example occurs between the phase of the hippocampal theta rhythm (5–10 Hz) and the γ amplitude in the hippocampus and neocortex^19,35^. As mentioned in the Introduction, the respiratory rhythm modulates γ activity in several areas of rodent and humans brain^11,19,23,29,30,32–34^. Interestingly, recent studies in humans show that the change from nasal to mouth breathing decreases CFC between theta and γ in the temporal lobe and alter limbic-based behavior^33^.

In the present study we show the existence of CFC between the phase of the respiratory wave and the amplitude of γ oscillation in the cat neocortex. It is important to note that the average frequency and limits of the γ burst differ between species^22, 32–34^, and varies according to the animal’s alertness level^32^. In addition, we show that this coupling remains intact during the carbachol-induced cataplexy, which shows that the muscular tone is not involved in this phenomenon.

The existence of CRP and CFC(Resp-γ) in *Rodentia, Primate* and our results in *Carnivora*, suggests a preserved mammal trait. Given the evolutionary importance of smell and the dense OB connectivity, it is highly probable to find similar phenomena in other non-mammalian vertebrates.

### Are CFC(Resp-γ) and inter-cortical γ coherence part of the same phenomenon?

Nasal breathing is part of “active sensation”, where the animals produce motor actions specifically tuned to obtain useful sensory information about their environment^32,39–41^. Examples are whisking and sniffing in rodents, electrolocation in fishes, echolocation in bats and odontocetes cetaceans as well as fingers and eye movements in primates^32,42,43^. In primates, the visual exploration is related to eye movement, specifically saccades and micro-saccades^44,45^, which are highly rhythmic^20,21^. In visual areas there are low frequency oscillations phase-locked to the rhythmic saccadic movements which exhibits a clear CFC with the γ band activity^20,21^. Something similar happens in the cortex with whisker movement in rats^32,39^ and also with nasal respiration^19,23,29,32,38^. Furthermore, slow waves coupled to saccadic movement are capable of modulating inter-cortical spikes and LFP γ coherence in visual areas^20^. Recently, we showed that during REM sleep, hippocampal theta activity modulates the coherence of intra-hemispheric high frequency oscillations (110-160Hz) in medial and posterior cortices of the rat^18^. In addition, the present work shows, for the first time, that long-range γ coherence occurs modulated by the respiratory phase, suggesting a unified phenomenon. This “respiratory binding effect” is observed at the neocortical level; however, the OB γ activity seems not to be involved in this mechanism.

Recent works propose that respiratory rhythms facilitated inter-regional communication via CFC(Resp-γ) ^19,22^. Other lines of research propose that phase synchrony between areas of the brain, especially at γ frequencies, constitutes a dynamical mechanism for the control of cross-regional information flow^3,46,47^. The findings of this work can potentially bring together these two theoretical frameworks in a global neocortical processing scheme; i.e. high frequency interregional binding is modulated by physiological rhythms such as respiration.

### Cortical respiratory entrainment is not present during sleep

We demonstrated that the CRP, CFC(Resp-γ) and inter-cortical γ coherence are absent during NREM and REM sleep. Hence, breathing appears to be unable to entrain the oscillatory activity of the neocortex during sleep. Recently, Zhong *et al*.^19^ showed that during REM sleep no CRP or CFC(Resp-γ) is observed in any neocortical areas of the rat. Nevertheless, during NREM sleep they observed power spectrum peaks that coincide in frequency between the respiration and pre-limbic LFPs. On the other hand, they also observed CFC between slow cortical oscillations and γ activity (CFC_(slow-γ)_) in the OB and pre-limbic areas. However, two matching frequency peaks do not necessarily imply two coherent timeseries at that frequency. It is also important to note that during NREM sleep there are slow cortical waves in the same frequency range as breathing (delta, 0.5-4 Hz). In this sense, this delta activity may generate the CFC(slow-γ) observed in the OB and pre-limbic areas. In this regard, Manabe & Mori^10^ showed that during REM and NREM sleep breathing was unable to entrain γ activity in the OB^10^.

Cognitive activity and different electrographic rhythms are generated by the activity of cortical and sub cortical neurons, which are reciprocally connected. These networks are modulated by the activating or waking-promoting systems of the brainstem, hypothalamus and basal forebrain that directly or indirectly project to the thalamus and/or cortex^2,48^. By regulating thalamocortical activities, these activating systems produce electrographic and behavioral arousal. The activating systems decrease their activity during the NREM sleep. However, while most monoaminergic systems decrease their activity during REM sleep (REM-off neurons), cholinergic neurons increase their discharge during this behavioral state (REM-on neurons), which contributes to cortical activation^2,48^. In addition, because these cholinergic neurons are active during REM sleep, they should not be critical to the generation of CRP and CFC_(Resp-γ)_ which is absent during this state. In fact, systemic muscarinic antagonists do not block CRP and CFC_(Resp-γ)_ in the dorsal hippocampus^28^. More efforts must be made to unravel what neurotransmitters are involved in the generation and maintenance of this cortical respiratory entrainment.

## Conclusions

The results obtained in rodents, humans and the present results in felines strongly suggest that CRP and CFC(Resp-γ) is a conserved phenomenon in mammals. Extending previous findings to the cat, we showed evidence supporting the dependency on behavioral state of the neural-respiratory coupling. We also demonstrated for the first time, that nasal respiration can modulate inter-cortical coherence at γ frequency, especially between remote neocortical areas. This evidence suggests that the respiratory rhythm could facilitate inter-regional communication^19,22^. Previously described γ synchrony between areas of the brain as a dynamical mechanism for the control of cross-areal information flow or “communication through coherence”^3,46,47^, could be part of a larger phenomenon which includes respiratory modulations. The strong modulation of the electrocortical activity by the respiratory rhythms could be the foundation of the effect of breathing on critical functions such as cognition, affection and stress responses^34,49^.

## Materials and Methods Experimental animals

Six adult cats were used in this study. Part of these animals have been employed in previous studies^36^. The animals were obtained from and determined to be in good health by the Institutional Animal Care Facility. All experimental procedures were conducted in accordance with the *Guide for the Care and Use of Laboratory Animals* (8th edition, National Academy Press, Washington DC, 2011), and were approved by the Institutional Animal Care Commission (N^o^:070153-000089-17). Adequate measures were taken to minimize pain, discomfort or stress. In addition, all efforts were made to use the minimum number of animals necessary to produce reliable scientific data.

### Surgical procedures

The animals were chronically implanted with electrodes to monitor the states of sleep and W^2,9,36,37^. Prior to being anesthetized, each cat was premedicated with xylazine (2.2 mg/kg, i.m.), atropine (0.04 mg/kg, i.m.) and antibiotics (Tribrissen®, 30 mg/kg, i.m.). Anesthesia, which was initially induced with ketamine (15 mg/kg, i.m.), was maintained with a gas mixture of isoflourane in oxygen (1-3%). The head was positioned in a stereotaxic frame and the skull was exposed. Stainless steel screw electrodes (1mm diameter) were placed on the surface (above the dura matter) of different cortical areas (Figure S1). In addition, bipolar electrodes were implanted in both lateral geniculate nuclei (LGN) to monitor ponto-geniculo-occipital (PGO) waves, and in the orbital portion to record the electro-oculogram. The electrodes were connected to a Winchester plug, which together with two plastic tubes were bonded to the skull with acrylic cement, to maintain the animals in a stereotaxic head-fixed position without pain or pressure. In all animals, a craniotomy was drilled in the skull overlying the cerebellar cortex, filled with bone-wax and was subsequently used to provide access to the pons for carbachol administration^36,37^. After the animals had recovered from the preceding surgical procedures, they were adapted to the recording environment for a period of at least two weeks^9,36^.

### Experimental sessions

Sessions of 4 hours were conducted between 11 A.M. and 3 P.M in a temperature-controlled environment (21-23°C). During these sessions (as well as during the adaptation), the animals’ head was held in a stereotaxic position by four steel bars that were placed into the chronically implanted plastic tubes, while the body rested in a sleeping bag^9,36^.

The ECoG activity was recorded with a monopolar (referential) configuration, utilizing a common reference electrode located in the left frontal sinus^9,36^. Controls experiments were made using other references electrodes^9^. Bipolar electromyogram (EMG) of the nuchal muscles was also monitored. The electrocardiogram, by electrodes acutely placed on the skin over the pre-cordial region, and respiratory activity by means of a micro-effort piezo crystal infant sensor and a thermistor located in the nostril were also recorded. Each cat was recorded daily for approximately 30 days to obtain complete data sets.

Bioelectric signals were amplified (x1000), filtered (0.1-500 Hz), sampled (2048 Hz, 2^16^ bits) and stored in a PC using the Spike 2 software (CED). Data were obtained during spontaneously occurring W, NREM, REM sleep, and during the induction of REM carbachol (REMc) or cataplexy (CA)^36,37^.

### Carbachol microinjection into the NPO

To induce REMc or CA, carbachol (0.8 µg in 0.2 µL of saline) was microinjected unilaterally for a period of 1 minute into the NPO with a Hamilton syringe^36,37^. Carbachol microinjections were performed either during NREM sleep or W. Two successful carbachol microinjections (i.e. in these experiments REMc and CA episodes were generated) were carried out for each animal (Figure S1; cats 3-6). The eyes of the animals were examined throughout the recording sessions to determine if they were closed or open, and if the pupils were mydriatic or miotic. We also monitored the degree of relaxation of the nictitating membrane and whether the animals were able to track visual or auditory stimuli^36,37^. During CA, the ECoG resembles W, PGO waves in the LGN were not observed, the eyes were open with moderate pupillary dilatation and auditory and visual stimuli were tracked as during natural W. In contrast, during REMc, the ECoG and PGO waves in the LGN did not differ from naturally-occurring REM sleep (Figure 5; arrowheads). Additionally, the eyes were closed, and the nictitating membrane was relaxed^36,37^. REMc and CA share the same muscular atony (Figure 5A, EMG).

### Data analysis

Sleep and W were quantified in 10 second epochs applying standard classification criteria^9,36^. Then, the maximum number of non-transitional and artifact-free periods of 30 s was selected for analysis during each behavioral state^18^. For each animal, we analyzed up to four complete recording to guarantee a minimum 500 seconds length for each cat and behavioral state (REM sleep is the limiting factor since it is a small percentage of the total recording time). This data was imported and analyzed offline using built-in and custom-written MATLAB codes (Mathworks). All data was previously filtered (low-pas 100 Hz) and down sample at 256 Hz to decrease the computational load of subsequent analyzes.

**Power Spectrum** was calculated by means of Welch’s periodogram (built-in MATLAB *pwelch* function).

**Coherence spectra** (Figures 1B,5B and 6E) of electrode pairs were computed using magnitude-squared coherence (built-in MATLAB *mscohere* function) and Fisher z’ transform was applied. Both power and coherence spectra calculations were carried out in all the data segments using 10 seconds Hamming windows and a frequency resolution of 0.1 Hz.

**CCF-map** (Figures 3 and S3) was generated between respiration and the envelopes of frequencies between 20-100 Hz (custom-written MATLAB code). To obtain the CCF-map, several band-pass filtered signals were generated from the raw recordings (built-in MATLAB *eegfilt* function). We used 10 Hz bandwidth and 5 Hz steps, covering from 15 up to 105 Hz.

The CCF-map was then generated by means of a raster plot of CCFs (built-in MATLAB *xcross* function) calculated between respiration and the envelopes (built-in MATLAB *hilbert* function) of each filtered signal.

**PAC and MI** (Figures 4 and 5D) were calculated used the framework previously described by Tort *et al*.^35^. First, the phase of the respiratory wave was extracted (built-in MATLAB *hilbert* function). Second, the γ band was band pass filtered (30-60 Hz) and envelopes were generated (*eegfilt* and *Hilbert*, respectively). Phase-amplitude plots were computed using 20° phase bins of the respiratory signal. The mean amplitude in each phase bin was normalized by the sum across bins, so that amplitude values in each plot summed to 1. MI=(Hmax-Hpac)/Hmax, where Hmax is the maximum entropy (Shannon entropy) value that can be obtained from the phase-amplitude relations (uniform distribution) and Hpac is the entropy of the phase-amplitude relations for the original signal^35^. MI→1 means maximum PAC while MI→0 means absence of PAC. **γ Coherence in function of the respiratory phase** (Figure 6C) was calculated (custom-written MATLAB code) using the phase of the respiratory signal and the coherence between pairs of the ECoG signals. After the respiratory phase extraction (*hilbert*) the pair of ECoG recordings was divided into nine parts (40° each) taking as reference the respiratory wave. Then the coherence of each bin phase was computed using magnitude-squared coherence (*mscohere*). Then, the spectral coherence was plotted as a function of the respiratory phase as a heat map.

**Phase looking value (PLV) as a function of the respiratory phase** (Figures 6D and S4) were calculated (custom-written MATLAB code) using the bin phase of the respiratory signal and the phase coherence (PLV) between each pair of the ECoG signals. After the respiratory phase bin extraction (*hilbert*) each pair of ECoG records was band-pass filtered (10 Hz bandwidth and 5 Hz steps, covering from 15 up to 105 Hz; or 30-60 Hz) and phase extracted (*eegfilt* and *hilbert*, respectively). Then, the ECoG phase difference was computed and the mean phase difference in the complex plane of each respiratory phase bin (20°) was calculated (PLV)^50^. PLV→1 means that phase difference is constant through all the respiratory bin and PLV→0 means that phase difference changes randomly through respiratory bins.

**R^2^ in function of the respiratory phase** was calculated (custom-written MATLAB code) using phase bin of the respiratory signal and the square root of the linear correlation between a pair of electrodes. After the respiratory phase bin extraction (*hilbert*) each pair of ECoG recordings was band-pas filtered (30-60 Hz; *eegfilt*) and then square root of the linear correlation (R^2^; built-in MATLAB *corr* function) was calculated with a moving window of 0.088 seconds (four γ cycles) with ≈99% of superposition. Then, the mean of R^2^ was calculated for each respiratory phase bin (Figure S5B). We also calculated the coherence between the γ R^2^ wave and the respiratory waveform (Figure S5A). **Statistics**. Group data are expressed as mean ± standard deviation. Most of the statistical analyses were assessed by paired two-tailed t-test (see Results and Figures’ legends). The significance of the differences among behavioral states was evaluated with repeated measures ANOVA (rmANOVA) along with Bonferroni *post hoc* tests. When sphericity criteria were not accomplished (tested by Mauchly’s test), the Greenhouse-Geisser correction was applied. The criterion used to reject null hypotheses was p<0.05. For Figures 6C and S4, a paired two-tailed t-test was performed with a Bonferroni correction for multiple comparisons. With this correction, p<0.0001 was considered statistically significant.

## Acknowledgements

This study was partially supported by the ‘Programa de Desarrollo de Ciencias Basicas, PEDECIBA’, ‘Comision Sectorial de Investigacion Cientifica’, CSIC. And Agencia Nacional de Investigación en Innovaciónn (ANII)’. S. CZ. and N. V. received a postgraduate scholarship from the ANII, while M. C. received a fellowship from ‘Comisiónn de Apoyo a Postgrados (CAP)’ D.RL. acknowledges funding from Proyecto Fondecyt 3120185.

## Authors Contributions

Financial support. P.T. Experimental design. M.C., S.CZ., P.T. Experimental procedures. M.C., S.CZ., N.V., P.T. Analysis of the data. M.C., J.G., D.RL., N.R. Discussion and interpretation of the data. M.C., P.T., J.G., D.RL., N.R., S.CZ., N.V. Wrote the manuscript. M.C., P.T. All the authors participated in critical revision the manuscript, added important intellectual content, and approved the final version.

## Data Availability

For access to data and custom computer code contact Dr. Pablo Torterolo and Matías Cavelli.

## Abbreviations

γ: gamma
ACF: autocorrelation function
ANOVA: ANalysis Of VAriance
CA: cataplexy
CCF: cross-correlation function
CFC: cross-frequency coupling
CRP: cortical respiratory potential
EEG: electroencephalogram
ECoG: electrocorticogram
EMG: electromyogram
i.p.: intraperitoneal
L: left
LGN: lateral geniculate nucleus
LFP: local field potential
M1: primary motor cortex
MI: modulation index
NPO: nucleus pontis oralis
NREM: non-REM sleep
REMc: REM carbachol
OB: olfactory bulb
OSN: olfactory sensory neurons
PAC: phase-amplitude coupling
Pf: prefrontal cortex
PGO: ponto geniculo occipital
PLV: phase locking value
REM: rapid eyes movement
rmANOVA: repeated measures ANOVA
RMS: root mean square
s.c.: subcutaneous
S1: primary somato-sensory cortex
SD: standard deviation
V1: primary visual cortex
V2: secondary visual cortex
W: wakefulness

